# Insects and incest: sib-mating tolerance in natural populations of a parasitoid wasp

**DOI:** 10.1101/169268

**Authors:** Marie Collet, Isabelle Amat, Sandrine Sauzet, Alexandra Auguste, Xavier Fauvergue, Laurence Mouton, Emmanuel Desouhant

## Abstract

1. Sib-mating avoidance is a pervasive behaviour that likely evolves in species subject to inbreeding depression. Laboratory studies have provided elegant demonstrations of sib-mating avoidance, but small-scale bioassays often minimize the costs associated with mate finding and choice, which could lead to spurious findings.
2. We used the hymenopteran parasitoid wasp *Venturia canescens* as a model organism, because previous laboratory studies revealed that sib-mating led to a 25% decrease in fertile offspring, and that sib-mating was partially avoided.
3. Our study consisted of a mate choice experiment in laboratory cages to determine if kin discrimination occurs in this species. We further performed a field study in which 86 wild-caught males, 155 wild-caught females and their 226 daughters daughters were genotyped at eighteen microsatellite loci. With these field data, we reconstructed the genotype of each female’s mate and estimated the relatedness of each mating pair.
4. Mate choice experiments confirmed that females are capable of discriminating kin. Time to mating depended on the frequency of female encounters with related and unrelated males. Contrary to previously published results, however, no sib-mating avoidance was detected. In the field, the effective rate of sib-mating did not differ from the probability that sibs encounter one other at random, which corroborates the absence of sib-mating avoidance. We also detected a weak but significant male bias in dispersal, which could reduce encounters between sibs.
5. Our results suggest that, despite kin discrimination, *V. canescens* tolerates sib-mating in the field. The weak male-biased dispersal cannot explain entirely this pattern. This raises the question as to why kin discrimination is maintained in this species. It further calls into question the idea that inbreeding depression occurs in most species with single-locus complementary sex determination.

## Introduction

When inbreeding lowers fitness-related components, such as survival and fertility, natural selection should favour behaviours preventing the reproduction of genetically-related individuals, or mitigating harmful consequences referred to as inbreeding depression (Pusey and Wolf, 1996; Keller and Waller, 2002; Angeloni et al., 2011). Inbreeding depression is caused by the expression of deleterious recessive alleles at the homozygous state or by a loss of heterosis in inbred crosses (Charlesworth and Charlesworth, 1987; Pusey and Wolf, 1996). Selection on behaviours underlying inbreeding avoidance should thus depend on inbreeding load, which balances the advantages of inbreeding avoidance with the costs of implementing adapted physiological or behavioural responses. Selection may also depend on the benefits of inbreeding in terms of inclusive fitness, as inbreeding results in an increased representation of genes identical by descent in the offspring (Kokko and Ots, 2006; Puurtinen, 2011; Duthie and Reid, 2016).

Responses to inbreeding risk reflect the benefits and costs associated with inbreeding and inbreeding avoidance. Observations range from systematic inbreeding avoidance to inbreeding tolerance or even inbreeding preference (Szulkin et al., 2013). For example, inbreeding preference was observed in the cichlid fish *Pelvicachromis taeniatus*, where inbreeding does not the lower survival and growth rate of young (Thünken et al., 2007). Related pairs further invest more in parental care and cooperate better. In rodents, such as *Marmota flavipentris*, inbreeding is preferred despite lower survival of inbred progeny. This preference can be explained by the large variation of male reproductive success and the composition of social groups, where males mating with related females do not suffer decreased annual reproductive success compared to the majority of males with little or no reproductive success (Olson et al., 2012). In contrast, mole rats or ring-tailed lemurs living in socially structured groups avoid inbreeding. In mole rats, avoidance leads to a high reproductive skew in males, with a subset of males that never reproduce (Cooney and Bennett, 2000). In ring-tailed lemurs, inbreeding avoidance allows females to save energy (i.e. invested in pregnancy and lactation) that would have be allocated to the production of inbred offspring, which suffer from depressed immune functions and have a reduced life expectancy (Boulet et al., 2009; Charpentier et al., 2008). It is, therefore, essential to quantify the costs and benefits of inbreeding avoidance to understand the consequences of inbreeding and the evolution of avoidance behaviours.

Several behavioural and physiological strategies to avoid inbreeding risks have evolved in animals, including sib-mating avoidance, dispersal (Greenwood et al., 1978; Szulkin and Sheldon, 2008), polyandry, extra-pair paternity, divorce (Hatchwell et al., 2000; Cornell and Tregenza, 2007; Cohas et al., 2008; Lardy et al., 2011; Reid et al., 2015; Duthie and Reid, 2016), and postcopulatory mechanisms, such as the preferential use of sperm from unrelated males by inseminated females (Tregenza and Wedell, 2002; Bretman et al., 2004). Mate choice is probably the most pervasive behaviour for sib-mating avoidance (Pusey and Wolf, 1996) and relies mostly on sib recognition mediated by chemical communication (Howard and Blomquist, 2005; Johansson and Jones, 2007; Charpentier et al., 2008a; Bonadonna and Sanz-Aguilar, 2012). It is nonetheless important to note that kin recognition mechanisms can arise from selective forces other than inbreeding avoidance, such as nest-mate recognition, parental care, territory defence, and dominance hierarchies among others (Mateo, 2004). The balance between the costs and benefits associated with inbreeding avoidance also depend on environmental factors, such as population density, spatial or social population structure. A low population density constrains mate availability and encounter rates and may, therefore, influence mate choice (Kokko and Rankin, 2006; Duthie and Reid, 2016). In the lizard *Zootoca vivipara*, for example, female choosiness is reduced when mate encounter rates decrease (Breedveld and Fitze, 2015). Moreover, in the marsupial carnivore, *Antechinus agilis*, habitat selectivity following natal dispersal is negatively correlated with the abundance and relatedness of females occupying the novel habitat suggesting a pervasive effect of inbreeding risk on dispersal (Banks and Lindenmayer, 2014). In cooperatively breeding species, inbreeding avoidance strategies can lead to the suppression of reproduction outside dominant pairs due to the lack of opportunity for outbred matings (as in the Damaraland mole-rats *Fukomys damarensis*; (Cooney and Bennett, 2000). Reproductively supressed subordinates that are related to the dominant pair should, therefore, increase their involvement in raising offspring to increase their inclusive fitness and maintain the social structure of the group (Nichols, 2017). The importance of environmental factors in shaping inbreeding avoidance thus requires field data to be able to quantify costs and benefits associated with each strategy.

Assessing inbreeding avoidance patterns, that is, the occurrence of inbreeding avoidance and behavioural strategies implied, is a difficult task that requires the estimation of relatedness coefficients between actual and potential mates (Szulkin et al., 2013). This may explain why most field studies have been conducted on large species of mammals and birds, for which monitoring is much easier compared to small invertebrates (Cohas et al., 2008; Herfindal et al., 2014; Arct et al., 2015; Hardouin et al., 2015). It is thus not surprising that inbreeding avoidance patterns have been rarely documented in insects in the wild (but see Bretman et al., 2011 and Robinson et al., 2012). Laboratory studies nonetheless suggest various strategies for inbreeding avoidance in different insect genera, such as precopulatory avoidance of related males in butterflies (Fischer et al., 2015) and postcopulatory choice, where stored sperm from unrelated males is used preferentially by females of the cricket *Gryllus bimaculatus* (Bretman et al., 2004, 2009). In a social species of termite, *Neotermes chilensis*, dispersal of colony founding individuals is the main mechanism to avoid inbreeding (Aguilera-Olivares et al., 2015). In contrast, absence of inbreeding avoidance has also been documented in insects, including parasitoid wasps (Bourdais and Hance, 2009) and *Drosophila melanogaster* (Tan et al., 2012). The diversity of strategies unveiled in laboratory experiments calls for a more thorough investigation of inbreeding avoidance patterns under natural conditions, particularly in insects.

Here, we take up this challenge by studying a parasitoid wasp with a simple form of inbreeding depression, both in the laboratory and in the field. Parasitoids are haplodiploid, where males develop from unfertilized haploid eggs and females develop from fertilized diploid eggs. Many hymenopteran species further have a single-locus Complementary Sex Determination (sl-CSD) mechanism, with which sex is determined by both ploidy and heterozygosity at a unique sex determination locus (Cook, 1993; van Wilgenburg et al., 2006; Heimpel and de Boer, 2008; Asplen et al., 2009; Schmieder et al., 2012). Inbreeding depression arises for diploids that are homozygous at the sex locus, because inbreeding leads to a proportionally higher number of diploid males that are generally unviable or sterile (Cook, 1993; van Wilgenburg et al., 2006; Fauvergue et al., 2015). Whatever the diversity of sex alleles in the population, full sibs have 50% chance of sharing a common sex allele and, if they mate, half of their diploid offspring will develop into unfit males (Cook, 1993). Assuming females fertilize half of their eggs, sib-mating thus results in 12.5% fewer offspring, on average, in monandrous females such as *V. canescens*. sl-CSD is thus a form of inbreeding depression underlined by overdominance, with no deleterious alleles at the sex locus, but a strong heterozygous advantage (e.g., Charlesworth and Willis, 2009). The null fitness of diploid males should, therefore, favour sib-mating avoidance in species with sl-CSD.

The parasitoid wasp *Venturia canescens* Gravenhorst (Hymenoptera: Ichneumonidae) has a single locus complementary sex determination (Beukeboom, 2001) and inbreeding reduces the fitness of both males and females through the production of sterile diploid offspring. Inbreeding further has a negative impact on egg load and hatching rate (Vayssade et al., 2014; Chuine et al., 2015). In no-choice bioassays, mating success consistently decreased with increasing genetic relatedness between mates (Metzger et al., 2010a; Chuine, 2014). There is some evidence suggesting that females are the choosy sex, which makes sense in a species with a monandrous/polygenous mating system (Metzger et al., 2010a). When females were confronted with a choice between a brother and an unrelated male, however, females did not avoid inbreeding, potentially due to the small size of test tubes (i.e. where mixing of chemical signals may have occurred; Metzger et al., 2010a). In this study, we first tested the effect of male density and average relatedness on inbreeding avoidance in the laboratory. We expected sib-mating avoidance to decrease with decreasing density and increasing relatedness. We further implemented a population genetic approach based on microsatellite genotyping to assess inbreeding avoidance and underlying behaviours in field populations of *V. canescens*. For this, we sampled two natural populations of *V. canescens* and compared the observed rate of sib-mating to the probability of random encounters among genetically-related individuals. Under the hypothesis of sib-mating avoidance, we expected the rate of sib-mating to be lower than the probability that relatives encounter each other at random. We further used these field data to test whether sex-biased dispersal reduces encounters among sibs in the field. We show that sib-mating tolerance occurs in the wild and propose an evolutionary scenario under which sib tolerance may evolve without loss of the ability to discriminate kin. We discuss why our results differ from earlier findings of inbreeding avoidance in this species, as well as prior expectations about inbreeding depression in species with sl-CSD (van Wilgenburg et al., 2006).

## Materials and Methods

### Biological model

*Venturia canescens* is a monandrous solitary endoparasitoid found in most circum-Mediterranean areas (Salt, 1976). Females lay eggs in a large range of hosts found on various fruits (Salt, 1976; Driessen and Bernstein, 1999). Knowledge of the mating system of *V. canescens* under natural conditions is limited, but can partly be inferred from results of laboratory experiments. Females do not discriminate between host patches with more than four host larvae, suggesting that, in the field, females rarely encounter aggregated host (Driessen and Bernstein, 1999). It is, therefore, unlikely that *V. canescens* experiences local mate competition, where males remain at the natal patch to mate (Hamilton, 1967). Males search for females via a synergistic combination of volatile sex pheromones emitted by females and kairomones released by hosts (Metzger et al., 2010b). Both immature development and adult life lasts around three weeks (at 25°C), and females lay eggs during most of their adult life (Metzger et al., 2008), leading to overlapping generations. Depending on the study considered, sex-ratio is either considered balanced (Beukeboom, 2001) or weakly biased toward females (Metzger et al., 2008). Equal sex ratios are indeed expected when hosts do not aggregate (Driessen and Bernstein, 1999), local mate competition does not occur (Macke et al., 2012), and mate-finding takes place via volatile cues. The proportion of diploid males further corresponds to that expected when sex ratios are equal (Fauvergue et al., 2015). Field and laboratory studies further revealed that adult females are good dispersers, with a flight velocity estimated at 0.2 m.s^−1^ (Schneider et al., 2002; Desouhant et al 2003; Amat et al., 2012).

### Insect rearing

For laboratory experiments, we used a standardized laboratory rearing that had been established with about 60 females collected in the field near Valence (same as the current location), southern France, on several occasions during the summer of 2015. The host *E. kuehniella* was reared in the laboratory on organic wheat semolina medium for three weeks before being exposed to parasitoids (eggs were obtained from the Biotop rearing facility located in Livron sur Drôme, France). Parasitoid development took place in a controlled environment (temperature: 24±1°C; relative humidity: 60±5%; photoperiod: LD 12:12).

Experiments on female mate choice required related and unrelated males. We thus initiated families by collecting females randomly from the mass rearing (from different rearing boxes), and maintaining them individually in a circular box (Ø: 8 cm, height: 3 cm) with a meshed lid containing two males and about 30 three week-old host larvae on 1 cm of organic wheat semolina. The wasps were fed *ad libitum* with 50% water-diluted honey and allowed to lay eggs during their whole life. Approximately three weeks later, the parents were removed from the box if still alive and offspring collected at emergence, before reproduction. Daughters were kept individually in plastic vials, and males were kept with their brothers, both with access to a drop of water-diluted honey. All individuals used in the experiments were two days old.

### Insect sampling in the field

Adult *V. canescens* were captured from June to September 2014 during 10 non-consecutive days. Two field sites, located 300 km apart, were sampled: Valence (N 44°58′21″ E 4°55′39″, composed of fruit trees in an orchard) and Nice (N 43°41′23″ E 7°18′6″, composed of carob trees in a semi-natural park). We captured only females in Valence and wasps of both sexes in Nice.

Two different types of traps were used: kairomone-baited traps for the capture of live females, and yellow sticky traps for males (which were generally found dead). Female traps constituted of an open cup containing host larvae, *Ephestia kuehniella* Zeller (Lepidoptera: Pyralidae), along with rearing medium (semolina). Saliva secreted by host larvae when feeding serves as a kairomone for female *V. canescens* (Corbet, 1971). Traps were hung in trees (Metzger et al., 2008) and visited every 20 minutes to collect attracted females. For males, 125 mm × 200 mm yellow sticky cards were baited with extracts from females and hosts, a method that already proved successful previously (Collet et al., 2016). For each trap, a vial containing the extracts was attached to the centre of the card. For females, traps were hung within trees and all *V. canescens* males that were found stuck on the traps collected and conserved individually in 96% ethanol to preserve DNA. We captured 77 females and 86 males in Nice and 78 females in Valence (see Results and Table 3).

Females collected in the field were brought back to the laboratory, maintained in a climatic chamber under constant conditions (temperature: 24±1°C, relative humidity: 60±10%; photoperiod: L:D 16:8) and fed 50% water-diluted honey. Each female was allowed to lay eggs individually in *E. kuehniella* hosts for three days. As we could not control for the age and egg-laying experience of captured females, the number of offspring was highly variable (1-10 daughters and 1-7 sons). As sexual (arrhenotokous) and asexual (thelytokous) strains coexist in *V. canescens* (Beukeboom et al., 1999; Schneider et al., 2002), we used the presence of males among offspring as evidence for arrhenotoky, and thelytokous individuals were discarded. Mothers and their offspring were then preserved individually in 96% ethanol.

### Effect of male density and sib frequency on sib-mating avoidance in the laboratory

We aimed to untangle the effects of male density and the frequency of encounters with related males on female mate choice. To test this, we designed a factorial experiment with two densities of males (high density, noted *D*, with nine males, and low density, noted *d*, with three males) and two frequencies of related males (high frequency, noted *F*, with 2/3 of brothers, and low frequency, noted *f*, with 1/3 of brothers). Males were marked according to kinship (brother vs unrelated) with a drop of paint applied on their thorax 24h before being tested. The two colours used do not modify behaviours (Desouhant et al., 2003), but were nonetheless alternated randomly across trials. Each trial took place as follows: a virgin female was released in a mesh cage (30 × 30 × 30cm) containing males at a temperature of 25°C, a relative humidity of 65% and under constant luminosity. Courtship and mating behaviours were monitored by visual observation for 30 min or until a mating had occur between 10 am and 4 pm. We recorded mating latency, copulation duration, relatedness of the successful male, and courtship effort *(i.e*. the time and the duration of each male courtship behaviour) (van Santen and Schneider 2002). For each courtship sequence, we identified the relatedness (colour) of the courter; hence all courtships were counted independently of mating success. Twenty trials were performed per factor level, except for *d-F* (18 replicates).

We analysed the proportion of sib-mating with a Generalized Estimating Equation – Generalized Linear Model (GEE-GLM; Liang and Zeger, 1986, GEEPACK package in R software) implemented with a binomial error distribution, a logit link function, an exchangeable correlation structure, and a fixed scale. The model was fitted with mate relatedness as a response variable (binomial: 0 if the female mated with her brother, 1 if she mated with an unrelated male), density and frequency of brothers as main effects and in interaction as explanatory variables, and families as a random factor *(i.e*. clusters, Højsgaard et al., 2006). We added an offset term to take into account the theoretical mating proportion with related males *(i.e*. 1/3 in low frequency *f* and 2/3 in high frequency *F*). We further tested the effects of density and proportion of brothers on male courtship effort and latency before copulation. For courtship effort, we implemented a GLM based on a binomial distribution and a logit link function, with the percentage of courtships done by brothers as a response variable and the density and proportion of brothers as explanatory variables. For latency before copulation, we performed a Cox model with the same explanatory variables as fixed effects and family as a random effect (COXME package, Therneau, 2015 in R software). We also included the number and timing of courtship (from a related or unrelated male) as a time-dependent variable. If females assess relatedness with males during courtship, an increasing number of encounters with related males could lead the female to decrease her selectivity and, as a consequence, the latency before mating. Alternatively, an increasing number of encounters with unrelated males should not affect her selectivity and therefore her latency to mate.

### Genotyping

Genotyping was performed at 18 microsatellite loci developed by Collet et al. (2016) and Mateo Leach et al. (2012) (See SEM-A for details on the genotyping and SEM-D for details on the number of microsatellites used). We genotyped all field-captured males and females (hereafter referred to as wild females) and up to three daughters per female (see SEM-B for a justification of the number of offspring genotyped). When fewer than three daughters emerged (ca. 34% of which 40% produced only one daughter) all were genotyped.

We estimated polymorphism and linkage disequilibrium at each microsatellite locus for the wild females of Nice and Valence with the GENEPOP software version 4.3 (Raymond and Rousset, 1995). The frequency of null alleles was estimated with FREE NA software (Chapuis and Estoup, 2007). population differentiation was estimated based on Wright’s *F* estimator *F_st_* with the FSTAT software version 2.9.3 (Goudet, 1995). Effective population sizes were estimated using the full likelihood method, assuming random mating, developed by Wang (2009), and implemented in the software COLONY (Jones and Wang, 2010). As effective population sizes were small (< 80), we did not check for Hardy-Weinberg equilibrium. We also determined the number of diploid males as a proxy for consanguinity and the number of alleles at the *csd* locus. A male was considered diploid if at least one microsatellite locus was heterozygous, haploid otherwise (Collet et al., 2016).

### Inference of relatedness among mates

#### •Male genotypes

We inferred the genotype of wild females’ mates based on the genotypes of wild females and their daughters. For this, we used the software COLONY, which reconstructs families based on haplodiploid genotypes (Jones and Wang, 2010). When alternative genotypes were proposed for a given male, we selected the one that was compatible with the genotype of the daughter and the mother. When several alternatives were possible (e.g. if mother and daughters were heterozygous for the same alleles at a given locus), it was impossible to assign the right allele to the father with certitude. In this case, we assigned the allele with the highest probability of occurrence (the probability is given by COLONY. We successfully inferred the genotype of 51 mates for the 54 wild females captured (see Results and Table 3).

#### •Relatedness between mates

To assess if *V. canescens* females avoid mating with relatives in the field, we compared the observed number of wild females mated with a related male to the number expected under the assumption of random mating. For this, we determined a “relatedness threshold” above which individuals were considered genetically related. To determine this threshold, we simulated populations with similar allele frequencies at microsatellite loci to that of natural populations, and a balanced sex ratio (see SEM-C for details on simulations). Simulations allowed to keep track of pedigrees, yielding a perfect knowledge of kinship for all potential mates, which we classified in three categories: full-sibs, half-sibs, or unrelated. Using microsatellite genotypes from the same simulated dataset, we estimated the relatedness coefficient (r) for all female-male pairs (TrioML coefficient, software COANCESTRY; Wang, 2011). We then used these estimated relatedness coefficients with different relatedness thresholds to assign each dyad to related vs unrelated categories (SEM-C). By comparing these posterior assignations to the pedigrees known a priori, we estimated the relatedness threshold that minimized the number of wrong assignations (*i.e*. unrelated pairs assigned related based on the threshold, or vice-versa) under the random encounter hypothesis. We found *r_crit_* = 0.16 (SEM-D, Fig. S2). Logically, this threshold is lower than the theoretical relatedness between a brother and a sister (*r* _sister-brother_ = 0.25; *r* _brother-sister_ = 0.5) and in the interval of relatedness for 1^st^ degree cousins (*r* = 0.125 or *r* = 0.1875 depending on the cross). With this threshold, we expect to wrongly assign 11.4% of the relationships.

We applied this *r*_crit_ = 0.16 threshold to field data. First, we estimated the number of wild females that had been inseminated by a related male. Second, we generated a theoretical distribution of the number of couples with an r ≥ 0.16 under the null hypothesis of random mating. For this, each of the 51 wild females from Nice, for which a mate genotype had been successfully inferred, was paired randomly with one of the 79 genotyped males, and we counted the number of pairs with r ≥ 0.16. This calculation was repeated 1000 times to produce the null distribution. We finally compared the observed number of related pairs with the null distribution via a permutation test implemented in the function AS.RANDTEST from the ADE4 package in R software version 3.2.3 (Chessel et al., 2004; R Core Team, 2015). We also estimated the probability that a female encountered a related male in the field the day of her capture. Indeed, partners present on the same day in the same area have a high chance of encountering each other and mate due to the high dispersal capacity of this species.

### Sex-biased dispersal in the field

Sex-biased dispersal can decrease the burden of inbreeding depression by reducing encounters and subsequent mating among sibs. We thus assessed the dispersal ability of males and females from our genotyping data in the population of Nice, where two trapping locations were distinguishable (Figure 1). The first patch contained a group of 21 trees and the second patch 15 trees. Trees were located a few meters from one another within patches, and the two patches were about 200 m away. Six sampled trees further apart were not included in this scheme (black dots in Figure 1).

**Figure 1.**
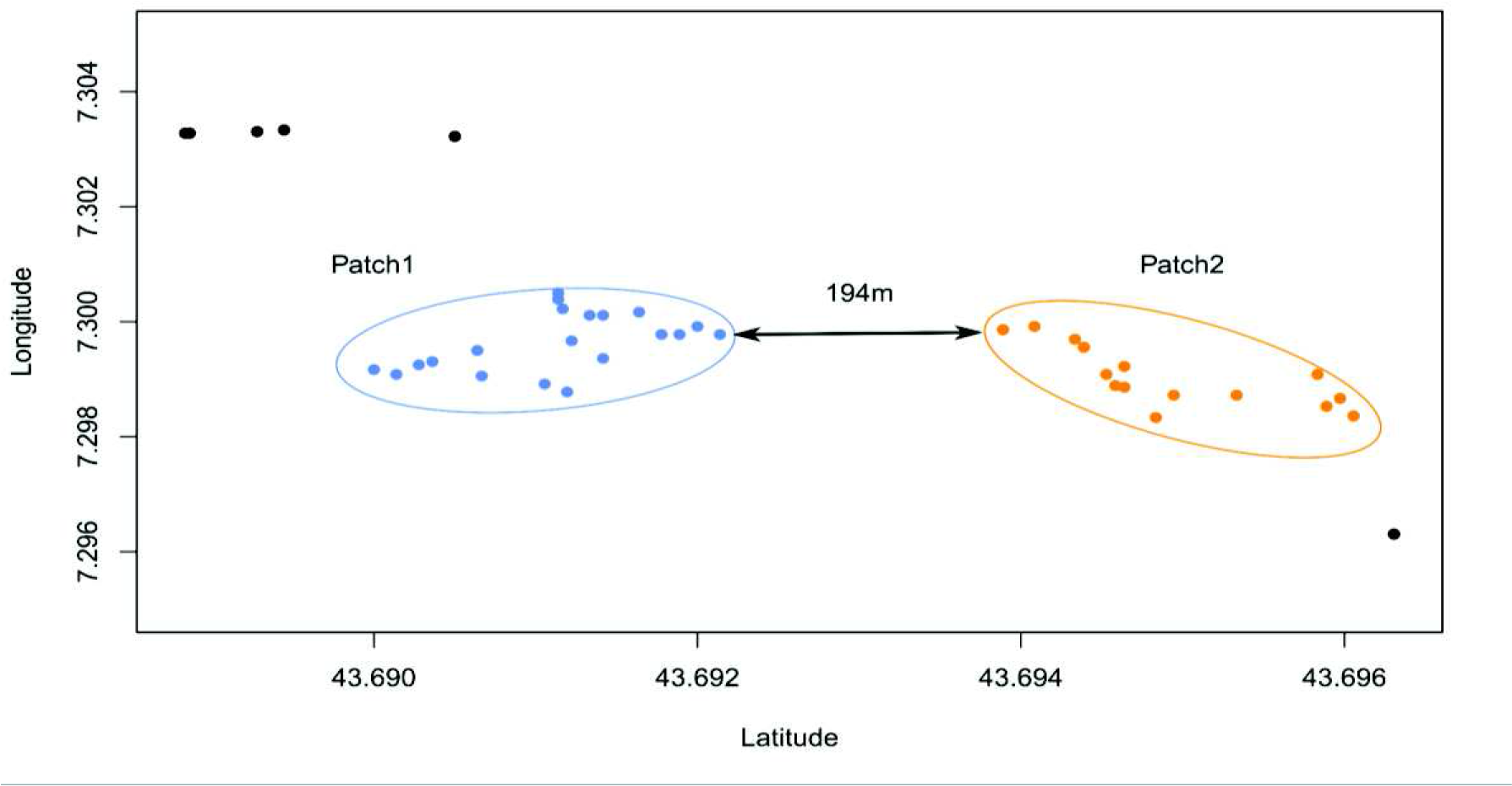
Location of carob trees at the Nice field site where *V. canescens* were captured during the summer of 2014. Each dot represents a tree. Axes are GPS coordinates in decimal degrees. Blue dots represent trees of “Patch 1” and orange dots represent trees of “Patch 2” (see also Table 4). Six trees (black dots) were not included because they fell outside the main patch areas.

We estimated the relatedness between individuals of each sex within and between patches using COANCESTRY software (Wang, 2011). Sex-biased dispersal should result in higher coefficients of relatedness within patches than between patches for the sex dispersing at smaller distances (Prugnolle and de Meeus, 2002). Coefficients of relatedness inter-patch and intra-patch were compared via a Kruskal-Wallis test (with patch as an explanatory factor) for each sex separately. When significant, post-hoc Mann-Whitney’s *U*-tests (with Bonferroni corrections) between the three modalities (*i.e*. relatedness within patch 1, relatedness within patch 2, and relatedness between patches) were performed.

### Results

#### Effect of male density and sib frequency on sib-mating avoidance in the laboratory

Mating success was higher at high male density (*D*; 81.6%, 40 matings/49 trials) than at low density (*d*; 44.1%; 38/86) (Table 1; proportion test: *χ*^2^ = 16.44, *df* = 1, *p* = 5.02 .10^−5^). The observed proportion of sib-mating did not depart from that expected under random encounters (GEE-GLM model, p-value of the intercept = 0.3). This was confirmed by a positive relationship between the proportion of related males and the probability of sib-mating (Wald statistic table, *χ*^2^ = 15.36, *df* = 1, *p* = 8.9 .10^−5^). Male density had no effect on mate choice, neither as a main effect (*χ*^2^ = 0.92, *df* = 1, p = 0.34) nor in interaction with the proportion of related males (*χ*^2^ = 0, *df* = 1, *p* = 1). Males did not differ in their courtship effort, as the proportion of courtships observed for unrelated males and brothers did not depart from the expected values (GLM model, p-value of the Intercept = 0.16): for example, in the *D-F* modality with 6 brothers and 3 unrelated males, we expected that 2/3 of the courtships would be done by brothers (i.e. 13.3 on 20 matings) and observed 12 sib-matings. Time before copulation decreased with increasing male density and increasing number of sib rejections (Table 2), but it was not affected by the proportion of related males or by the number of rejections of unrelated males (Table 2).

**Table 1.** Effect of male density and sib frequency on sibmating avoidance in *V. canescens*. Two densities of males (*D*: high density = 9 males, and *d*: low density = 3 males) and two frequencies of brothers (*F*: high frequency = 2/3 brothers and *f*: low frequency = 1/3 brothers) were manipulated in a cage experiment. Presented are the number of successful matings for each factor level (20, except for the *d-F* modality that contained only 18 replicates.

| | | $D$ | $d$ |
| --- | --- | --- | --- |
| $F$ | Brother | 12 | 12 |
|  | Unrelated | 8 | 6 |
|  | Total | 20/23 trials | 18/44 trials |
| $f$ | Brother | 6 | 6 |
|  | Unrelated | 14 | 14 |
|  | Total | 20/26 trials | 20/42 trials |

**Table 2.** Effect of male density, sib frequency (and their interaction) and the number of male rejections on mating latency in *V. canescens*. We analysed the deviance table of the mixed-effect Cox model with the female family as a random effect. Significant effects are in bold.

| | Log likelihood | $\chi^2$ | df | p-value |
| --- | --- | --- | --- | --- |
| NULL | -351.51 |  |  |  |
| **Male density** | **-338.44** | **26.14** | **1** | **$3.17 \cdot 10^{-7}$** |
| Sib frequency | -337.57 | 1.74 | 1 | 0.19 |
| **Rejection of sib** | **-332.45** | **10.25** | **1** | **$1.36 \cdot 10^{-3}$** |
| Rejection of unrelated | -331.46 | 1.97 | 1 | 0.16 |
| Density × sib frequency | -331.46 | 0.01 | 1 | 0.92 |

#### Genotyping

A total of 241 wasps were captured in Valence and Nice. In Valence, 78 wild females were captured (Table 3) of which 35 produced a total of 86 daughters (Table 3; 5.1 ± 0.4 offspring, including 2.4 ± 0.2 daughters per female). In Nice, 86 males and 77 females were captured, of which 54 produced 140 daughters (Table 3; 4.8 ± 0.3 offspring, including 2.6 ± 0.1 daughters per female). We genotyped 467 of these individuals at 18 microsatellite markers. We obtained the genotype at all 18 microsatellite loci for 402 individuals, at 15-17 loci for 50 individuals, 1-10 loci for 11 individuals, and no amplification was obtained for 4 individuals (Table 3). Only individuals with successful amplification at >14 loci were included in the dataset, which represented 452 individuals (96.8%).

**Table 3.** Number of adult *V. canescens* captured from wild populations in Nice and Valence. Pop = population; Number of mothers = number of wild females captured in the field that produced at least one daughter; F = female, M = male.

| Pop | Sex | Wild individuals | Wild individuals successfully genotyped | Number of mothers | Total number of daughters | Daughters successfully genotyped |
| --- | --- | --- | --- | --- | --- | --- |
| Nice | F | 77 | 77 (100%) | 54 | 140 | 134 (96%) |
|  | M | 86 | 79 (92%) | - | - | - |
| Valence | F | 78 | 77 (99%) | 35 | 86 | 85 (99%) |

No linkage disequilibrium was detected and null allele frequency was always lower than 6.5% in the two populations. We found no genetic differentiation between the two populations (*F_st_* = 0.01). We, nonetheless, treated the two populations separately because the probability that a male captured in one population had inseminated a female captured 300 km apart is negligible. Effective population sizes were estimated to be 79 for Nice (IC 95%: [44; 82]) and 51 for Valence (IC 95%: [26; 56]). These values are low and not significantly different.

#### Sib-mating avoidance in the field? Relatedness of actual versus potential mates

In Nice, the mean coefficient of relatedness between wild males and females was *r* = 0.047 ± 0.001. We estimated that 11% ± 0.79% (IC 95%) of the relatedness coefficients between these individuals were higher than the threshold *r_crit_* = 0.16 (671 comparisons over 6083, from 79 males × 77 females); the corresponding pairs were thus considered genetically related. The average probability that a female encountered a related male in the field on the day of capture was 12.0% ± 4.9 (IC 95%) (Fig. 2).

**Figure 2.**
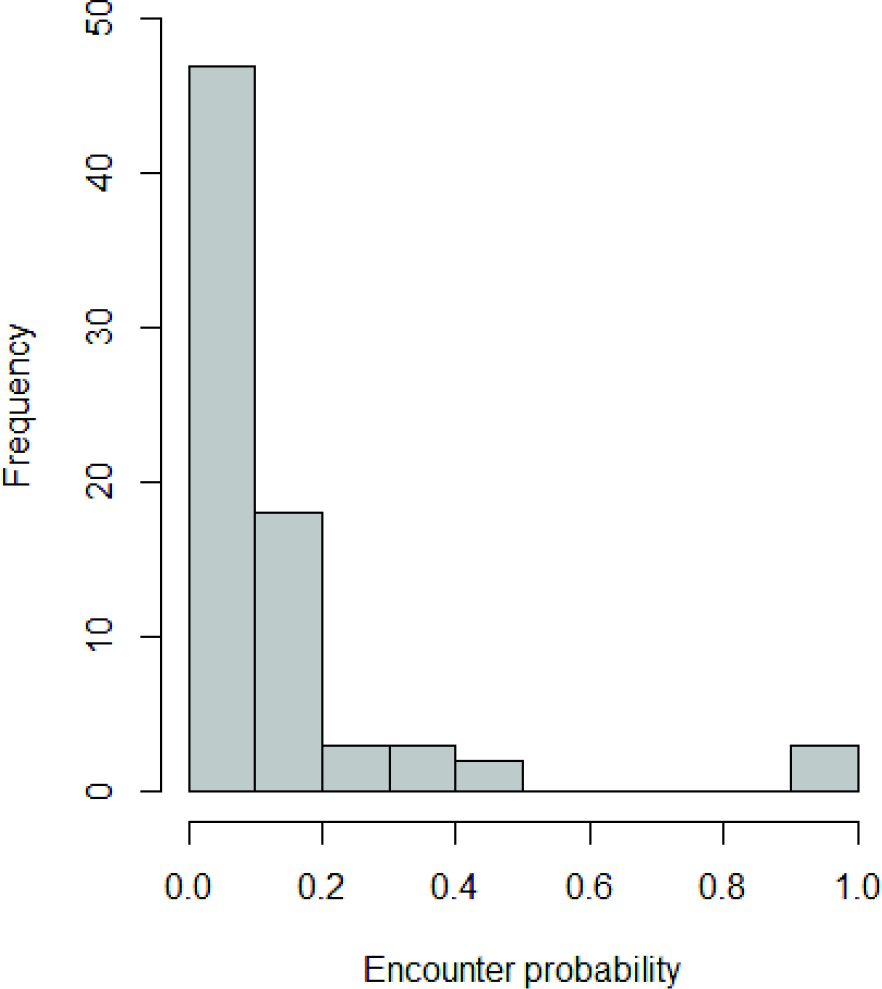
Distribution of the probability that a female *V. canescens* encountered a genetically related male during the day of capture in the population from Nice. For each female caught we computed her relatedness with all males captured the same day. We then counted the number of relationships with r ≥ 0.16 and divided this number by the total number of relatedness computed (*i.e.* the number of males captured the same day).

Thirty-five and 54 wild females from Valence and Nice, respectively, produced at least one daughter which allowed to infer the genotype of their mate (Table 3). The mean relatedness between these inferred pairs was 0.050 ± 0.009 in Nice and 0.044 ± 0.013 in Valence. Assuming *r_crit_* = 0.16, we found six and four genetically related pairs in Nice and Valence, respectively, that is, 11.1% ± 8.4% (IC 95%) and 11.4% ± 10.5% (IC 95%) of all pairs. For the population of Nice, this observed number of related pairs did not differ from the number expected under the assumption of random mating (1000 permutations; mean number of related pairs expected from random mating = 5.264; number observed = 6, *p* = 0.822; Fig. 3).

**Figure 3.**
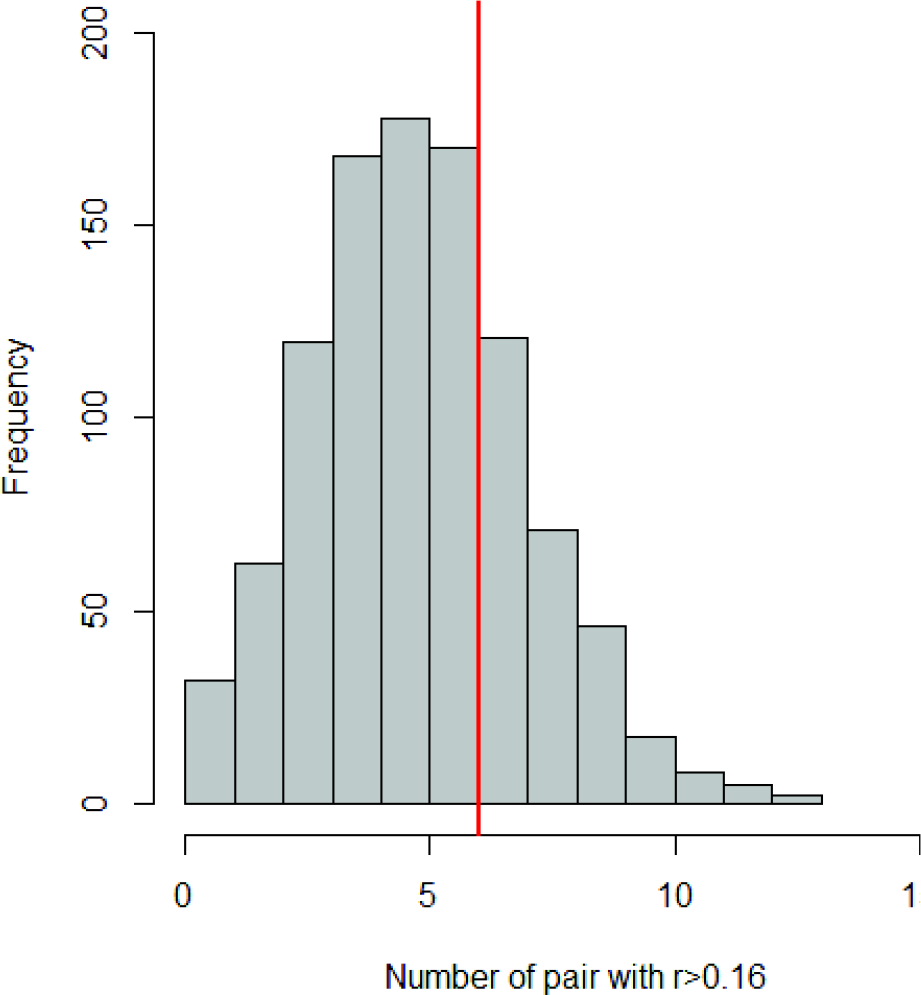
Theoretical distribution of the number of crosses between genetically related *V. canescens* males and females under the assumption of random mating. The theoretical distribution was built using random sampling of a mate for each of the 51 females from the Nice population, estimating r for each pair and counting those with r > 0.16. Sampling was reiterated 1000 times. The red line represents the observed number of pairs with a r > 0.16.

#### Proportion of diploid males and number of CSD alleles

We assessed the cost of inbreeding by estimating the number of diploid males in the population of Nice (no males were captured in Valence). In species with sl-CSD, diploid males result from matched matings (parents sharing a common allele at the *csd* locus), and the probability of matched mating in a panmictic population is 2/*k*, where *k* is the effective number of *csd* alleles. The number of alleles at the *csd* locus *k* could be estimated from the proportion of diploid males among the males *m* and the proportion fertilizations *s* (see SEM-E for the calculation). As previous findings suggest a balanced sex-ratio (Beukeboom, 2001; Fauvergue et al., 2015), we set s=0.5. We estimated the number of *csd* alleles in the population of Nice, where random matings is expected (see previous section). Six diploid males were found among the 79 males captured (*m*=7.6% ± 5.8%, IC 95%) yielding an estimation of 8.8 alleles at the *csd* locus and a probability 2/*k* = 0.23 of matched matings.

#### Sex-biased dispersal in the field

During the field experiment 50 females and 47 males were genotyped in patch 1, 18 females and 22 males in patch 2. Genetic relatedness was estimated among females and among males within patches (referred to as 1-1 for intra-patch 1 and 2-2 for intra-patch 2), as well as between patches (referred to as 1-2) (Table 4). Mean relatedness differed within and between patches for females (Kruskal-Wallis test: *χ*^2^ = 14.88, *df* = 2, *p* < 1.10^−4^), but not for males (Kruskal-Wallis test: *χ*^2^ = 1.72, *df* = 2, *p* = 0.424). For females, relatedness was similar within each of two patches (1-1 and 2-2; Mann-Whitney test: *U* = 80550, *p* = 0.520; Table 4) and higher than between patches (1-1 / 1-2 comparison, Mann-Whitney test: *U* = 56724, *p* < 4.10^−4^; 2-2 / 1-2 comparison, Mann-Whitney test: *U* = 65356, *p* = 0.012). Sex-biased dispersal, therefore, occurs in *V. canescens*, with males dispersing more than females.

**Table 4.** Mean relatedness r (± SE) among females and among males of *V. canescens*, either within patches (1-1 or 2-2) or between patches (1-2) of host plants (carob trees) in the population of Nice. F = female, M = male.

| Sex | Patch | Mean $r$ (± SE) |
| --- | --- | --- |
| F | 1-1 | 0.065 ± 0.002 |
| | 2-2 | 0.060 ± 0.006 |
| | 1-2 | 0.050 ± 0.002 |
| M | 1-1 | 0.042 ± 0.002 |
| | 2-2 | 0.048 ± 0.006 |
| | 1-2 | 0.043 ± 0.003 |

## Discussion

Sib-mating avoidance in *Venturia canescens* was previously observed in no-choice behavioural experiments conducted under laboratory conditions (Metzger et al., 2010a, Chuine, 2014). We show here that when females can choose between related and unrelated males, sib-mating does occur. The density of males and the proportion of related individuals in the surroundings do not change the pattern of inbreeding avoidance, suggesting that relatedness is not a cue for mate choice. Females nonetheless appear to discriminate sibs from non-sibs, because mating latency decreases after rejection of a sib. The sib-mating tolerance that we observed in the laboratory was confirmed by our genetic approach with field populations. We showed that the observed frequency of sib-mating corresponds to the probability of sib-mating expected when sib encounters occur at random. Our field data further suggest there is sex-biased dispersal, but without yielding passive sib-mating avoidance. We will next propose an evolutionary scenario to explain the maintenance of kin recognition, and its apparent absence of use in the mate choice context, in the laboratory and in the field.

In laboratory cages the proportion of sib-mating perfectly matched the proportion of sibs present. Mating rates between relatives, be they inferred from genetic analyses or from a simulation model assuming random encounters, were both equal to 11%. These two results are in line with expectations in the absence of sib-mating avoidance in *V. canescens*, and brings into question past evidence of kin recognition and avoidance in the context both of host parasitization (Bernstein and Driessen, 1996; Marris et al., 1996) and mate choice (Metzger et al., 2010a; Chuine et al., 2015). In particular, no-choice bioassays previously revealed that female *V. canescens* mated twice more often with unrelated males than with their brothers, and were therefore capable of sib-mating avoidance (Metzger et al. 2010a). In the presence of volatiles from brothers, females also discriminated against unrelated males, suggesting that mate-choice could be underlined by volatile cues (Metzger et al., 2010a; Chuine et al., 2015). Our laboratory cage experiment also suggests that females are able to recognize kin and modify a component of their mating behaviour (latency before copulation), but without sib-mating avoidance.

The lack of evidence for kin discrimination in our field study could result from a misclassification of cousins or more distantly related individuals as full sibs. When genetic relatedness decreases to that of cousins (r ≤ 0.1875), sib-mating avoidance in *V. canescens* declines steeply (Chuine, 2014). Misclassifying cousins as brothers would thus, unsurprisingly, yield an absence of sib-mating avoidance. This is, however, unlikely. We classified pairs as sibs versus non-sibs based on a threshold, *r_crit_* = 0.16, that proved to minimize the probability of misclassification (about 10% of errors). Moreover, increasing this threshold to further restrict the definition of kinship did not change our conclusions, that is, the observed number of related pairs was always similar to that expected under random mating (*rcrit* = 0.2: 1000 permutations, 2 observed related pairs, 1.88 expected related pairs, *p* = 1; *r_crit_* = 0.25: 1000 permutations, 0 observed related pairs, 0.999 expected related pairs, *p* = 0.637). The hypothesis that poor kinship assignation yielded spurious results thus falls short of argument, and we conclude that the lack of evidence for sib-mating avoidance truly reflects a lack of sib-mating avoidance, also called sib-mating tolerance.

Sex-biased dispersal can also shape the pattern of inbreeding avoidance in the field (Gandon, 1999; Roze and Rousset, 2005). In insects, male-biased dispersal was found mostly in social species (Johnstone et al., 2012; López-Uribe et al., 2014; Vitikainen et al., 2015). In non-social insects, Downey et al., (2015) found male-biased dispersal in more than 10 species of bean beetle. This dispersal pattern is consistent with the kin competition avoidance hypothesis (local mate competition hypothesis) but not with the inbreeding avoidance hypothesis. In *V. canescens* from Nice, we found a slight male-biased dispersal at a local scale (Table 4). It is unlikely that local mate competition explains the evolution of male biased dispersal in *V. canescens*, raising the question of the inbreeding avoidance hypothesis. However, the relatedness values between and within patches (between which dispersal occurs) in both sexes was low (between 0.042 and 0.065, Table2) and no structuration at larger scale (between Valence and Nice population) was found. If male-biased dispersal indeed affected the structure of this population and encounter rates between relatives, it should therefore be a very weak effect.

Density-dependent female choosiness in *V. canescens*, as shown in other species (Kokko and Rankin, 2006; Duthie and Reid, 2016), could also explain the absence of sib-mating avoidance. Avoidance of sibs would arise in dense or aggregated populations with high mate encounter rates, but not in scarce populations where the cost of discrimination – remaining virgin – is higher. This was shown in a small isolated population of moose, where females accepted mates with higher relatedness during years when males were less available (Herfindal et al., 2014). A similar reasoning may also apply to highly inbred populations, where sib-mating avoidance would result in a high probability of not mating at all, a penalty that could exceed the costs of inbreeding depression (Kokko and Ots, 2006). In *V. canescens*, population density indeed affects mating success but not sib-mating probability. Moreover, the density of related males that we tested in the laboratory were respectively twice and four times higher than those found in the field, and still, no sib-mating avoidance was observed. In the field, the distribution of the sib-encounter probability is skewed towards zero, leading to more than half of *V. canescens* females having no chance to encounter a related male on a particular day, possibly because of the low number of individuals in the field (Fig. 2). The average probability for a female to encounter a relative is nonetheless within the same range as actual sib-matings (12%; Fig. 2). Kin recognition mechanisms could then have persisted, but not sib-mating avoidance. We therefore conclude that *V. canescens* does not avoid sib-mating, whatever the density of males and level of inbreeding.

Female kin discrimination against related males could also decay over time. Encountering a succession of low-quality males (*i.e*. genetically-related in our study) could decrease female choosiness, similar to what was demonstrated for a monandrous wolf spider (Stoffer and Uetz, 2016). We have shown in our laboratory experiment that even if females did not avoid sib-mating, the latency before copulation decreased significantly after having rejected a brother. This result confirms that *V. canescens* is capable of discriminating relatives in a context of mating (see also Metzger et al., 2010a; Chuine, 2014; Chuine et al., 2015) and suggests that the presence of relatives could affect mating by decreasing the latency before copulation, as expected under this hypothesis. More experiments are, however, needed to firmly test the evolution of female choosiness over time.

Tolerance to sib-mating, such as that observed in *Venturia canescens*, is often explained by low inbreeding depression or high costs of outbreeding (Kokko and Ots, 2006; Puurtinen, 2011). We found a low effective population size at both sample sites (Ne = 51 in Valence, Ne = 79 in Nice), within the same range as threatened bumblebees (Ellis et al., 2006) or Hine’s emerald dragonflies (Monroe and Britten, 2015). *V. canescens* could be prone to loss of genetic diversity, possibly amplified by inbreeding tolerance, which would in turn increase inbreeding load. Alleles at the *csd* locus, the major mechanism of inbreeding depression in *V. canescens*, are, however, under strong balancing selection and are thus expected to overcome genetic impoverishment, similar to observations in populations of the Asiatic honeybee (Gloag et al., 2016). Moreover, inbreeding in *V. canescens* only has a moderate negative impact on egg load and hatching rate, and no effect on other fitness components, such as mating probability or body size (Vayssade et al., 2014). Despite common beliefs, the genetic load resulting from sl-CSD might not be dramatic in *Venturia canescens*. In the population of Nice, we estimated 8.8 alleles at the *csd* locus, which is in the range of what was previously found for other species in the order Hymenoptera (8-86 alleles, mostly between 9 to 20; Cook and Crozier, 1995). Under random mating, this yields a probability of matched-matings of 23% (2/8.8). In case of brother-sister mating, the probability of matched matings increases to 50%. Assuming that females fertilize half of their eggs so that matched matings results in 25% diploid males, and that diploid males are sterile (Fauvergue et al. 2015), the expected reduction in the number of fertile offspring under random mating would be 5.8% (0.23 × 0.25). If sib-mating was perfectly avoided, the expected reduction in the number of fertile offspring would be 4.8% (0.19 × 0.25, see SEM-E for the calculation). Sib-mating avoidance would thus result in about 1% more fertile offspring in the populations we studied, a fitness gain that may appear insignificant. The relatively low probability of matched matings and the consequent benign inbreeding load may thus be insufficient to select for sib-mating avoidance. *V. canescens* may thus be an example of a species with inbreeding tolerance (Waser et al., 1986; Kokko and Ots, 2006).

The expression of sexual selection is dependent of the social environment in which the individuals interact, and the laboratory experiments therefore add biases on mate choice study (Miller and Svensson, 2014; Nieberding and Holveck, 2017). The differences between previous laboratory study and our results could highlight these differences in social environments and underline the necessity to confront laboratory and natural conditions data to estimate mate selectivity. Data from the field should be considered as the more relevant and should help to identify biases related to lab conditions. Discrepancies between sibmating avoidance revealed in previous research and our finding of inbreeding tolerance could moreover highlight the evolution of kin-recognition mechanisms for purposes other than sib-mating avoidance. This would have produced artefactual behaviours when observed in small arenas (Metzger et al., 2010a), but vanishing in larger cages and in the field. Evidence for active kin recognition is pervasive in various ecological contexts such as territory defence, reciprocal altruism, or dominance hierarchies (Mateo, 2004). In *V. canescens*, the evolution of kin recognition for the avoidance of superparasitism could explain the maintenance of kin recognition without sib-mating avoidance. Indeed, females are able to discriminate the self, their relatives and their conspecifics when laying eggs, and avoid already parasitized hosts. As only a single offspring can develop in a host, superparasitism has a high cost for female fitness. Moreover, the female’s inclusive fitness costs increase in the case of superparasitism of a host already parasitized by relatives. Consistently, the intensity of superparasitism avoidance increases with genetic relatedness to the first laying female(Hubbard et al., 1987; Marris et al., 1996; Amat et al., 2009). If kin recognition evolved in this context, kin discrimination in a situation of mate choice could be a by-product of this primary selection on kin recognition. We could therefore expect a relatively general molecular pathway, which would have been expressed in a small-scale mating bioassay: kin recognition during mating is probably mediated by semiochemicals from the insect cuticle (cuticular hydrocarbons, Metzger et al., 2010a; Chuine, 2014) and discrimination against superparasitized larvae is driven by weak volatile hydrocarbon signals left by females during oviposition (Fisher, 1961; Harrison et al., 1985). These mechanistic similarities point towards a unique pathway for kin recognition. Superparasitism avoidance could act as a strong selective pressure for the maintenance of kin recognition, because the cost of superparasitism is positively correlated with the relatedness among competing females. Further experiments are needed to test the idea that sib-mating avoidance observed under laboratory conditions is a byproduct of cognitive abilities that evolved in the context of intraspecific competition.

## Authors’ contributions

M.C, I.A., X.F., L.M. and E.D. designed the experiment and M.C., S.S and A.A collected the data. M.C., I.A., L.M. and E.D. designed and interpreted the analyses. M.C. conducted the analyses and assembled a first draft. M.C., I.A., X.F., L.M. and E.D. all contributed to revisions.

## Acknowledgments

We want to thank François Debias for his help in the field capture of insects, rearing and technical assistance, and Aurélie Blin for technical advices on genotyping. We thank Angèle, Anatole and Auguste Desouhant for their help in the field sampling. We are most also thankful to Caroline Nieberding, Bertanne Visser and an anonymous reviewer for their careful reviews and helpful comments that improved the manuscript. This work was partly funded thanks to Agence Nationale de la Recherche (“Sextinction” project, ANR-2010-BLAN-1717). We declare no conflict of interest.

## Data availability

Data available from Zenodo Repository: https://zenodo.org/record/1184074

R scripts for statistical analysis available in Supplementary Electronic Material F (SEM-F).

